# Membrane-mediated cooperativity of antigen receptors and CD2 adhesion receptors in CAR-T cell activation

**DOI:** 10.64898/2026.10.27.762129

**Authors:** Thomas R. Weikl, P. Anton van der Merwe, Omer Dushek

## Abstract

Chimeric antigen receptor (CAR)-T cells exhibit a major defect in antigen recognition, often requiring 100–1000-fold higher antigen densities for activation than the native T cell receptor (TCR). Recent work has traced this defect to the inefficient use of CD2/CD58 adhesion by CAR-T cells and demonstrated that matching the extracellular dimensions of CD2/CD58 and CAR/antigen complexes can dramatically enhance antigen sensitivity. Remarkably, modest nanometre-scale changes in CD2/CD58 dimensions produce orders-of-magnitude changes in antigen sensitivity, while having little effect on T cell activation at high antigen densities. Here, we present a general model for the length-dependent cooperativity of antigen receptors and CD2 adhesion receptors in CAR-T cell activation, combining statistical-physical theory with coarse-grained simulations. The model predicts that the antigen concentration required for activation increases sharply with the length mismatch between antigen-receptor and adhesion-receptor complexes until a critical mismatch is reached, at which the complexes segregate. Beyond this critical mismatch when complexes segregate, antigen sensitivity becomes largely independent of complex length, consistent with experimental observations. For small to moderate mismatches, the antigen concentration required for activation exhibits a Gaussian dependence on length mismatch with a characteristic width of only a few nanometres, explaining how nanometre-scale changes in CD2 length can alter antigen sensitivity by orders of magnitude. Fitting the model to measurements of CD2-length-dependent CAR-T cell activation further enables estimation of characteristic molecular length scales and concentrations of antigen-receptor and adhesion-receptor complexes. These results provide a quantitative physical framework connecting nanometre-scale organization of the cell–cell interface to CAR-T cell antigen sensitivity and highlight molecular dimensions as a design parameter for T cell-based immunotherapies.

**SIGNIFICANCE:** Receptor and ligand proteins anchored to apposing cell membranes can only bind at local membrane separations that are close to the length of the resulting receptor-ligand complexes. The interplay of membrane separation and complex formation in cell-cell adhesion zones leads to the cooperative binding of receptors and ligands with matching complex lengths and to the membrane-mediated repulsion and segregation of receptor-ligand complexes with substantially different lengths. We present a statistical-physical theory for the length-dependent cooperativity of antigen receptors and adhesion receptors in the activation of chimeric antigen receptor (CAR)-T cells by adhering antigen-presenting cells. Our theory helps to understand why matching the lengths of adhesion-receptor and antigen-receptor complexes with protein engineering can increase the antigen sensitivity of cancer-therapeutic CAR-T cells by several orders of magnitude.

## INTRODUCTION

T cells play a central role in adaptive immunity and are activated when their T cell receptors (TCRs) recognize foreign peptides presented by major histocompatibility complex (MHC) molecules on apposing cell surfaces (1). These antigenic peptides typically originate from viral or bacterial proteins and thereby signal infection. To redirect T cell immunity towards cancer cells, T cells have been engineered to express chimeric antigen receptors (CARs), which incorporate intracellular signalling motifs from the TCR-CD3 signalling machinery but possess distinct extracellular domains that recognize non-MHC surface molecules on cancer cells. CAR-T cells have transformed the treatment of haematological malignancies and are now being developed to target solid tumours (2, 3). However, whereas native TCRs can activate T cells in response to even a single foreign peptide–MHC (pMHC) antigen (4–6), synthetic CARs typically require 100–1000-fold higher antigen densities for activation (7–11). This reduced antigen sensitivity is thought to contribute to cancer relapse by allowing tumour cells with low surface antigen expression to escape CAR-T cell recognition (2, 12).

T cells employ several mechanisms to achieve the extraordinary antigen sensitivity of the native TCR, including adhesion mediated by CD2/CD58 complexes. As the TCR and pMHC are small molecules that project only ∼7.5 nm from their respective membranes, T cells must bring their membranes to within ∼15 nm of the target-cell membrane to enable TCR/pMHC binding. This membrane alignment can be mediated by the abundant CD2 receptor binding its broadly expressed ligand CD58, as the resulting CD2/CD58 complex spans the same ∼15 nm distance (Fig. 1A) (13, 14). We previously showed that CD2/CD58 adhesion can enhance the antigen sensitivity of the native TCR by more than 100-fold, yet has a negligible effect on CAR antigen sensitivity (10, 11). We hypothesized that this difference arises from a mismatch between the extracellular dimensions of CAR/antigen and CD2/CD58 complexes (Fig. 1B). Consistent with this hypothesis, elongating the extracellular domain of CD2 dramatically enhanced CAR antigen sensitivity (15). Moreover, because CARs differ in extracellular architecture, owing in part to hinges derived from different human proteins (Fig. 1C), distinct CD2 lengths were required to optimize the antigen sensitivity of different CARs targeting the same antigen. Remarkably, modest nanometre-scale changes in CD2 length produced dramatic 10–100-fold changes in antigen sensitivity, demonstrating that CAR-T cell activation is exquisitely sensitive to the molecular dimensions of CD2/CD58 adhesion complexes. Moreover, we found that CD2 length did not impact CAR-T cell activation at high antigen densities.

**Figure 1:**
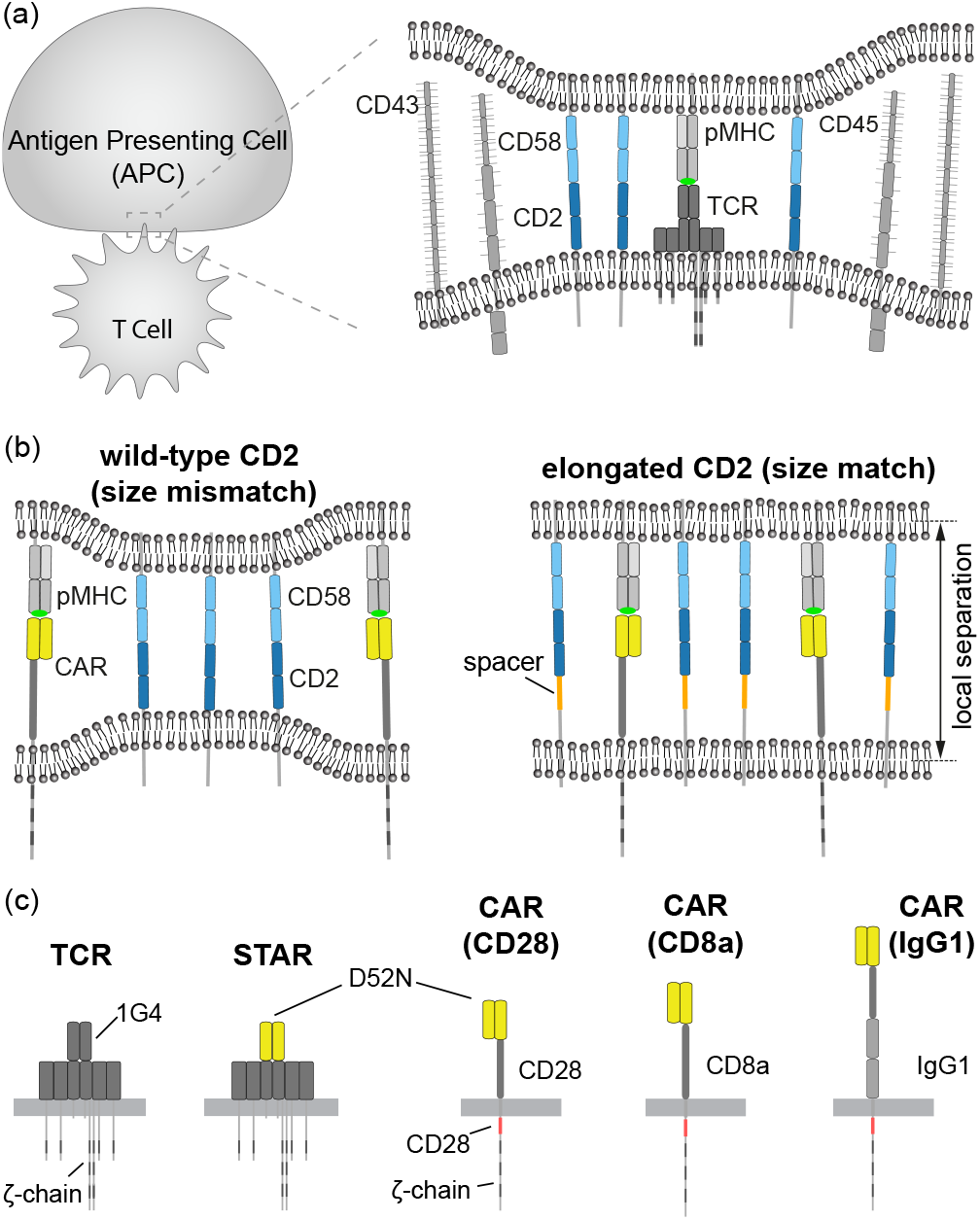
The CD2/CD58 adhesion complex is proposed to generate a ∼14 nm close-contact that efficiently enables the native TCR but not synthetic CARs to bind antigen. (a) T cells interact with antigen presenting cells (APCs) through close-contacts mediated by the CD2/CD58 adhesion complex that align membranes to the ∼14 nm distance required by the TCR to bind pMHC antigens. (b) Proposed size-based mechanism by which the wild-type CD2/CD58 adhesion complex cannot promote CAR/antigen binding (left) whereas a synthetic elongation of CD2 can promote CAR/antigen binding when size-matched (right) (15). (c) Schematics of the indicated antigen receptors, including the native T cell receptor (TCR) and 4 synthetic antigen receptors that all recognise the same NY-ESO-1 peptide antigen presented on HLA-A*02:01. The TCR relies on the 1G4 sequences whereas the synthetic antigen receptors rely on the D52N antibody. The STAR fuses the variable chains of the antibody to the TCR constant domains whereas the CARs rely on a single-chain variable fragment (scFv) of the D52N antibody fused to the indicated hinge and cytoplasmic domains. The affinity between the pMHC antigen and the TCR (K_D_ = 7.2 *µ*M), STAR (K_D_ = 1.3 *µ*M), and the scFv-containing CARs (K_D_ = 0.15 *µ*M) was previously measured (10, 16).

Here, we seek to quantitatively understand how the relative lengths of adhesion and antigen-receptor complexes determine antigen sensitivity in CAR-T cells. We develop a general, simulation-informed model of membrane-mediated cooperativity between antigen and adhesion receptors for small to moderate differences in complex length. The model enables characteristic molecular length scales and complex concentrations to be inferred from experimental measurements of CD2-length-dependent CAR-T cell activity (15). Consistent with experiments, our model and simulations show that the antigen concentration required for CAR-T cell activation increases sharply with increasing length mismatch between antigen-receptor and adhesion-receptor complexes, up to a critical mismatch at which the complexes segregate. Beyond this critical mismatch, the antigen concentration required for activation becomes largely independent of the mismatch, consistent with experimental observations that CARs of different extracellular dimensions can activate T cells at sufficiently high antigen densities. Together, these results provide a quantitative physical framework for understanding how nanometre-scale membrane organization controls receptor cooperativity and antigen sensitivity in CAR-T cells.

## METHODS

### General model for the cooperativity between adhesion receptors and antigen receptors in CAR-T cell activation

Our modeling of experimental data for the CD2-length-dependent activation of CAR-T cells is based on the equation (17)

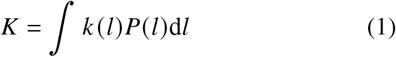

for the two-dimensional binding constant *K* of the antigen receptors, where *k*(*l*) is the binding constant as a function of the local membrane separation *l*, and *P*(*l*) is the distribution of the local membrane separation. The equation has been derived based on statistical-physical theory and coarse-grained simulations (17, 18) and reflects that the receptors can only bind to their antigen ligands if the local membrane separation *l* at the site of receptors and ligands is within an appropriate range, and that this local membrane separation varies along the membranes and in time because of thermally excited shape fluctuations of the apposing membranes, which are held together by adhesion and antigen-receptor/ligand complexes. In this modeling, we focus on the regime of small to moderate length mismatch between antigen-receptor complexes and adhesion-receptor complexes (CD2), and assume that the distribution *P*(*l*) of the local membrane separation is dominated by the much higher abundance of CD2 adhesion complexes, and essentially not affected by the very few antigen-receptor complexes required for activation in this regime. The distribution *P*(*l*) then can be approximated by the Gaussian function (17, 18)

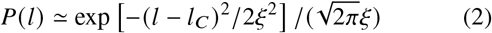

where *l*_*C*_ is the “preferred” local membrane separation for CD2 binding CD58, and the standard deviation ξ of the Gaussian function is the relative roughness of the apposing membranes.

We also approximate the separation-dependent binding constant *k* (*l*) of the antigen receptor by a Gaussian function

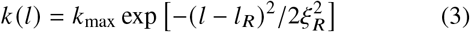

with maximum value *k*_max_, preferred membrane separation *l*_*R*_ for antigen-receptor binding, and standard deviation ξ_*R*_. The standard deviation ξ_*R*_ can be understood to reflect variations of the local separation at the site of a receptor/antigen complex that can arise from a tilting of the receptor complex and from the conformational flexibility of linker segments that connect the extracellular domains of the antigen receptor and the antigen ligand to their transmembrane domains (17, 19). Eq. (1) then leads to the expression

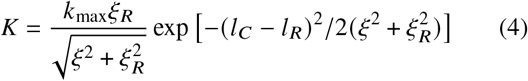

which quantifies how the two-dimensional receptor binding constant *K* depends on the length mismatch Δ*l* = |*l*_*C*_ − *l*_*R*_| between the CD2/CD58 adhesion complexes and the antigen-receptor complexes. According to eq. (1), an increase of the length mismatch Δ*l* causes a decrease of the receptor binding constant *K*. Because of the relation

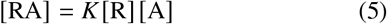

between the area concentrations RA, R, and A of the membrane-anchored antigen-receptor complex RA and the unbound receptor R and antigen ligand A, a decrease of the receptor binding constant *K* needs to be “compensated” by an increase in the antigen concentration A to reach a given receptor complex concentration RA . A common way to quantify antigen sensitivity in experimental work is to perform a peptide antigen titration and determine the concentration of the antigen that produces a half-maximal response (“EC50”). Within the context of the present model of receptor-ligand interactions at an interface, we assume that this EC50 is proportional to the concentration of membrane-presented antigen A that gives rise to a concentration of receptor-ligand complexes RA _50_ at which a half-maximal response is triggered. Based on eqs. (4) and (11), the antigen concentration required for a half-maximal response then can be expressed as

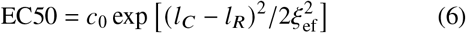

with 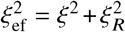 and a prefactor *c*_0_ that is independent of the length mismatch |*l*_*C*_ − *l*_*R*_| between the CD2/CD58 adhesion and antigen-receptor/antigen complexes.

### Coarse-grained model of CAR-T cell adhesion mediated by complexes formed by adhesion and/or antigen receptors

In our coarse-grained model of biomembrane adhesion, the membranes are represented as elastic surfaces, and the receptors and ligands as individual molecules that diffuse along the surfaces and bind if the local membrane separation at the site of an apposing receptor-ligand pair is within the binding distance for the specific receptor and ligand (20, 21). Such coarse-grained elastic models have been used extensively to simulate membrane adhesion via receptor-ligand complexes (20–33). The conformations of the two adhering membranes in these models are described in Monge parametrization by their local deviations out of a reference plane. In our model, this reference plane is discretized into a square lattice of quadratic patches *i* with side length *a*, which results in a discretization of the two membranes into apposing patches with local separation *l*_*i*_ (see Fig. 2(a)). The effective bending energy (34) of the two membranes then can be expressed as (20)

**Figure 2:**
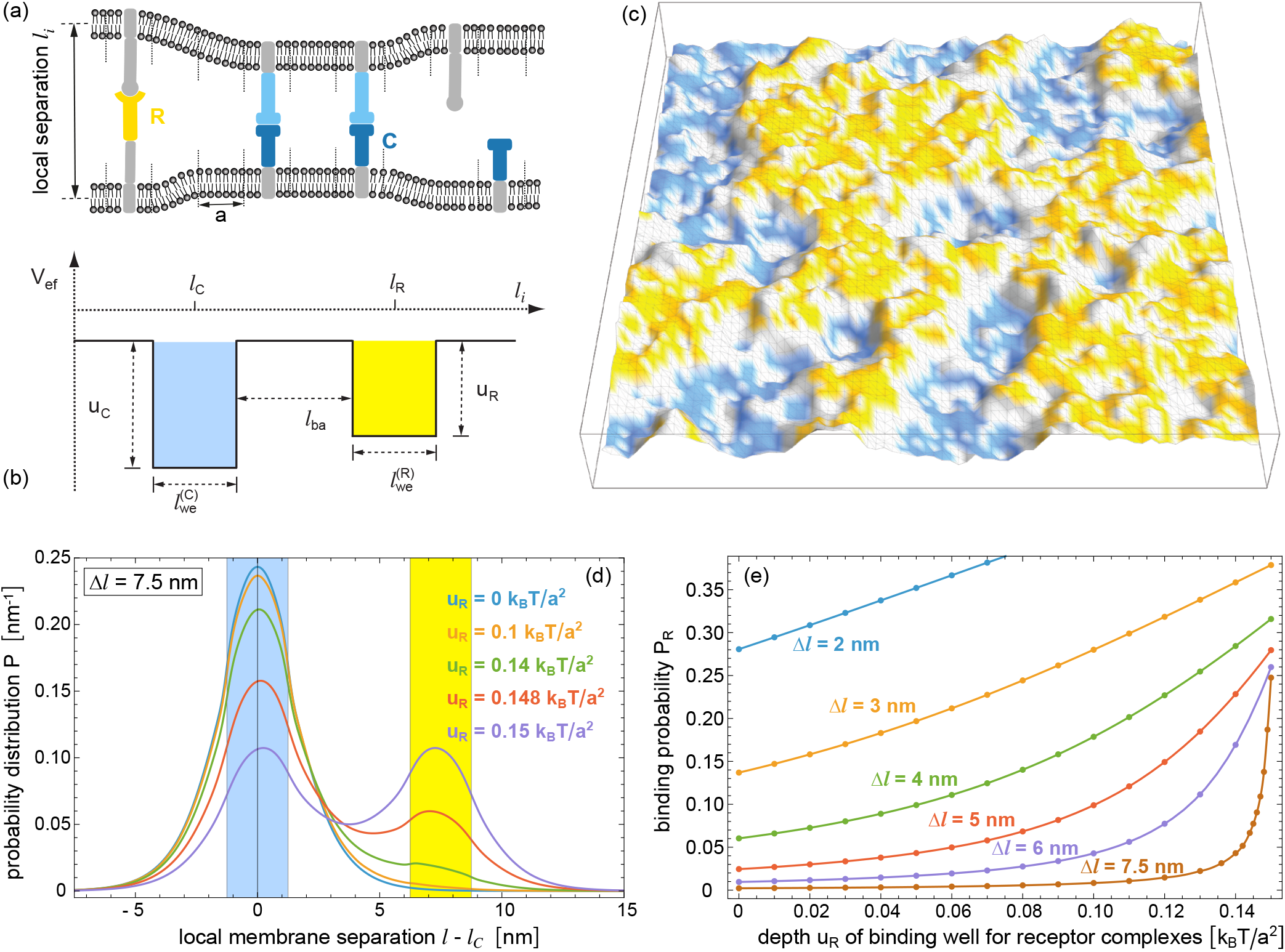
A coarse-grained spatial and stochastic model for biomembrane interactions quantifies the probability of antigen-receptor/antigen complexes when they have a length mismatch with adhesion-receptor/ligand complexes. (a) The coarse-grained model for biomembrane adhesion relies on discretizing the membrane in apposing quadratic patches with side length *a*, which can contain single receptor and ligand molecules. An apposing receptor-ligand pair can form a bound complex if the local separation *l*_*i*_ of their membrane patches deviates from the preferred values *l*_*R*_ and *l*_*C*_ of the antigen-receptor and the adhesion-receptor complex by less than 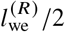 and 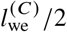, respectively. (b) Integrating over all possible distributions of the receptor and ligand molecules in this model leads to an effective double-well adhesion potential *V*_ef_ with wells of width 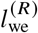 and 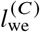 centered at *l*_*R*_ and *l*_*C*_ and separated by a “barrier” of width *l*_ba_. The well depths *u*_*R*_ and *u*_*C*_ depend on the concentrations and binding constants of the receptors and ligands (see Eqs. (9) and (10)). (c) Simulation shapshot of the separation field {*l*_*i*_} of two membranes interaction via *V*_ef_ for 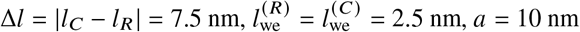, and *u*_*R*_ = *u*_*C*_ = 0.15*k*_*B*_*T*/*a*^2^. Patches with membrane separations *l*_*i*_ within the two binding wells are coloured in yellow and blue, respectively. (d) Probability distributions *P* (*l*) of the local membrane separation *l* obtained from Monte Carlo (MC) simulations. With increasing *u*_*R*_, the probability distribution *P* (*l*) is increasingly shifted towards the binding well for the antigen-receptor complexes and attains a symmetric bimodal shape for *u*_*R*_ = *u*_*C*_ = 0.15*k*_*B*_*T*/*a*^2^. (e) Probability *P*_*R*_ that the local separation *l* is within the binding well of the antigen-receptor complexes as a function of the well depth *u*_*R*_ for different values of the length mismatch Δ*l*. The data points represent results of MC simulations with a membrane size of 160 × 160 patches and a simulation length of 10^9^ MC steps per patch. The lines are interpolations of the data points. The statistical errors are smaller than the point sizes.

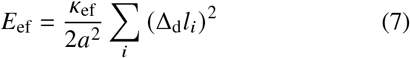

with effective bending rigidity *κ*_ef_, discretized mean curvature (Δ_*d*_*l*_*i*_)/*a*^2^, and discretized Laplacian Δ_*d*_*l*_*i*_ = *l*_*i*1_+*l*_*i*2_+*l*_*i*3_+*l*_*i*4_−4*l*_*i*_, where *l*_*i*1_ to *l*_*i*4_ are the membrane separations at the four nearest-neighbor sites of site *i*. The effective bending rigidity of the separation field {*l*_*i*_} of the two apposing membranes with rigidities *κ*_1_ and *κ*_2_ is (35, 36)

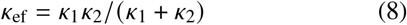

In our model, each membrane patch of the T cell membrane can contain a single antigen receptor (R) or single CD2 adhesion receptor (C), each membrane patch of the apposing cell membrane can contain a single antigen ligand (A) or single adhesion-receptor ligand (L), and the receptors interact with their ligands via square-well potentials. An antigen receptor R at membrane patch *i* of the T cell membrane binds to an antigen ligand A located at the apposing membrane patch *i* with binding energy *U*_*R*_ if the local separation *l*_*i*_ of the membrane patches deviates from the preferred binding separation *l*_*R*_ of the receptor complex by less than 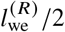, where 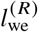 is the width of the binding well. Similarly, an adhesion receptor C at membrane patch *i* of the T cell membrane binds to an adhesion ligand L located at the opposing membrane patch *i* with binding energy *U*_*C*_ if the local separation *l*_*i*_ deviates from the preferred adhesion-receptor binding separation *l*_*C*_ by less than 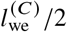.

Equilibrium properties of this model, such as the equilibrium concentration RA of receptor-antigen complexes, can be obtained from the partition function, which is the sum of all possible conformations Boltzmann-weighted by their energy, i.e. by the sum of the elastic energy of eq. (7) and the binding energies of all antigen and adhesion complexes of the conformation. Calculating the partition function thus requires to sum over all possible local separation fields *l*_*i*_ and all possible distributions of the receptor and ligand molecules on the membranes. In the grand-canonical ensemble in which the concentrations of the unbound receptors and ligands are adjusted by chemical potentials, the summation over all possible distributions of receptors and ligands can be carried out exactly, which leads to an effective model of homogeneous membranes interacting via an effective potential *V*_ef_ that depends on the concentrations and binding energies of the receptors and ligands (20, 21, 26). If the length difference | *l*_*C*_ − *l*_*R*_ | of the antigen and adhesion-receptor complexes is larger than 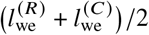, the effective potential *V*_ef_ (*l*_*i*_) is a double-well potential with well depths

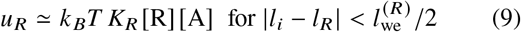

and

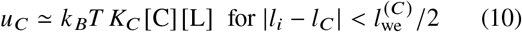

where *K*_*R*_ = *a*^2^ exp(*U*_*R*_/*k*_*B*_*T*) and *K*_*C*_ = *a*^2^ exp (*U*_*R*_/*k*_*B*_*T*) are the binding constants of the antigen receptor and adhesion-receptor complexes (see Fig. 2(b)). If the length difference |*l*_*C*_ −*l*_*R*_ | is smaller than 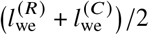, the effective potential *V*_ef_ (*l*_*i*_) adopts the value *u*_*R*_ + *u*_*C*_ for local separations *l*_*i*_ at which both complexes can form. Eqs. (9) and (10) can be derived from exact expressions for the well depths under the assumption that the fractions of membrane patches occupied by receptors and ligands is much smaller than the maximal value 1 at which each patch is occupied by a protein (26). In this effective model, the concentrations of complexes

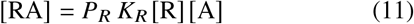

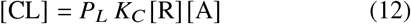

depend on the equilibrium probabilities *P*_*R*_ and *P*_*C*_ that the local separation *l*_*i*_ of the membranes is within the binding ranges 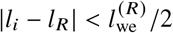 and 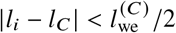, respectively (26).

## RESULTS

### A modest mismatch in extracellular length with adhesion complexes dramatically reduces antigen receptor binding to antigen

We first used our coarse-grained model of biomembrane adhesion to investigate the probability of antigen binding by antigen receptors, such as CARs, when they form complexes whose extracellular length is different from the length of adhesion-receptor complexes, such as CD2/CD58. The model allows to determine how the equilibrium concentration [RA] of antigen-receptor complexes depends on the antigen concentration [A], binding affinity, and the length difference *l*_*C*_ − *l*_*R*_ between the antigen-receptor and adhesion-receptor complexes. In this model, the apposing cell membranes of the T cell and APC are discretized into membrane patches that can contain single receptor and ligand molecules (see Fig. 2(a)). The shapes and shape fluctuations of the adhering membranes are reflected by the local separations of the apposing membrane patches, and the receptors and ligands diffuse along the membranes and can form complexes if the local separation *l*_*i*_ at the site *i* of an apposing receptor-ligand pair is within their binding distance (see Methods for details).

Determining equilibrium properties in this model with Monte Carlo (MC) simulations can be significantly simplified by first integrating out the degrees of freedom of the receptors and ligands in the model’s partition function, which leads to an effective model of two membranes interacting via a double-well potential *V*_ef_ with well depths *u*_*R*_ and *u*_*C*_ that depend on the concentrations and binding affinities of the receptors and ligands (see Fig. 2(b) and eqs. (9) and (10)). Besides the well depths *u*_*R*_ and *u*_*C*_, the parameters of this effective model are the widths 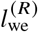 and 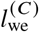 of the two binding wells, the effective rigidity *κ*_ef_ of the membranes given in eq. (8), the length difference *l*_*C*_ − *l*_*R*_ of the antigen and adhesion-receptor complexes, and the side length *a* of the discrete membrane patches. In this effective model, the well depth *u*_*R*_ is proportional to the concentration of unbound antigen [A] (see eq. (9)), and the concentration [RA] of antigen-receptor complexes is proportional to the probability *P*_*R*_ that the local separation *l*_*i*_ of the fluctuating membranes is within the binding range 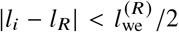 of the complexes (see eq. (11)). The remaining task of MC simulations then is to determine this probability *P*_*R*_ as central quantity for obtaining the antigen-receptor complex concentration [RA].

To illustrate our simulation approach, we first consider a situation with a length mismatch between the antigen-receptor and adhesion-receptor complexes of 7.5 nm (i.e. Δ*l* = |*l*_*C*_ − *l*_*R*_| = 7.5 nm). The MC simulations allow to determine the distribution *P l* of the local membrane separation *l* at various values of the well depth *u*_*R*_ (see Fig. 2(d)), which is proportional to the antigen concentration A and affinity. In these simulations, the bending rigidities of the two membranes have the typical value 20 *k*_*B*_*T* (37), which results in the effective rigidity *κ*_ef_ = 10 *k*_*B*_*T* (see eq. (8)), the widths of both wells are taken to be *l*_we_ = 2.5 nm, the well depth for adhesion complexes is *u*_*C*_ = 0.15 *k*_*B*_*T a*^2^, and the side length of the membrane patches is *a* = 10 nm. Without antigen binding (*u*_*R*_ = 0), the distribution *P l* is centered around the binding well of the adhesion complexes (see Fig. 2(d), blue region). The probability *P*_*C*_ that the local membrane separation is in the adhesion-complex well is about 0.55, which leads to the concentration [CL] = *P*_*C*_ *u*_*C*_/*k*_*B*_*T* ≃ 825/µm^2^ of adhesion complexes according to eqs. (10) and (12). As the affinity and/or concentration of antigen increases (i.e. as *u*_*R*_ increases), the probability distribution *P l* is shifted further and further towards the binding well for the antigen-receptor complexes and attains a bimodal shape with two equal maximal values for *u*_*R*_ = *u*_*C*_ = 0.15*k*_*B*_*T*/*a*^2^. The probability *P*_*R*_ that the local membrane separation is in the antigen-receptor well thus increases with *u*_*R*_.

To explore how the length mismatch Δ*l* affects antigen binding, we determined the binding probability *P*_*R*_ in the antigen-receptor well for different antigen concentrations/affinities (i.e. different values of *u*_*R*_) and for different values of Δ*l*. The binding probability increases with the antigen concentration/affinity (i.e. for increasing *u*_*R*_) but decreases with increasing length mismatch Δ*l*. As the length mismatch increases, we find that *P*_*R*_ *u*_*R*_ becomes steeper and steeper close to *u*_*R*_ = *u*_*C*_ = 0.15 *k*_*B*_*T*/*a*^2^, which indicates that the critical length difference Δ*l*^*^ for lateral segregation is approached. To determine Δ*l*^*^, we make use of the fact that the correlation length of the membrane shape fluctuations becomes infinite at the critical point of segregation and obtain Δ*l*^*^ = 8.4±0.1 nm from an analysis of MC results for different membrane sizes (see Appendix B).

To quantify how the antigen concentration [A] required for CAR-T cell activation depends on the length mismatch Δ*l* = |*l*_*C*_ − *l*_*R*_| of the antigen-receptor and adhesion-receptor complexes, we now assume that a half-maximal activation response occurs at a certain concentration [RA]_50_ of the antigen-receptor complexes. Because the antigen concentration A is proportional to *u*_*R*_, we numerically determine the value 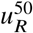 of the effective well depth *u*_*R*_, at which the concentration [RA]_50_ is achieved, from the equation

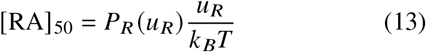

with functions *P*_*R*_ (*u*_*R*_) obtained from MC simulations (see Fig. 2(e)). This equation follows from eqs. (9) and (11). Fig. 3 illustrates how 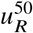 depends on the length difference *l*_*C*_ − *l*_*R*_ of the antigen-receptor and adhesion-receptor complexes for three different, exemplary values of RA _50_. The full data points are determined from eq. (13), and the open data points from the approximative equation

**Figure 3:**
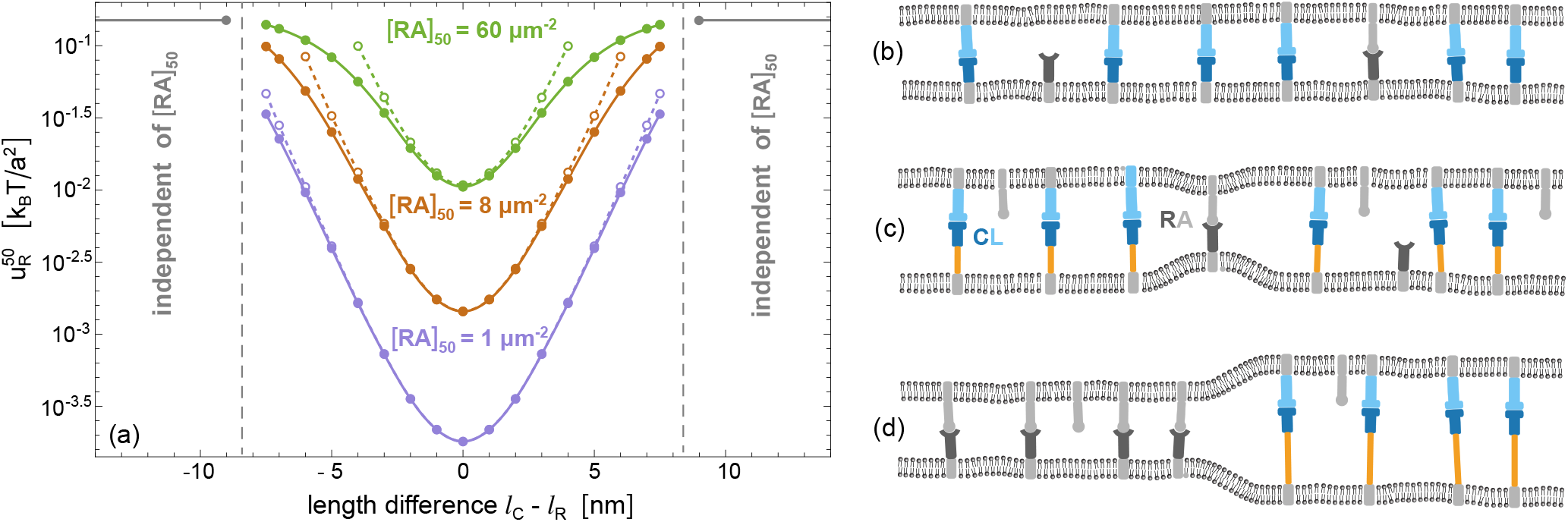
A modest extracellular length mismatch between adhesion and antigen-receptor complexes of <10 nm can reduce antigen binding by 100-fold. (a) Threshold value 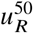 of the well depth *u*_*R*_ at which the complex concentration RA _50_ for half-maximal CAR-T cell activation is achieved as a function of the length difference *l*_*C*_ − *l*_*R*_ of the antigen-receptor and adhesion-receptor complexes, for three different exemplary values of RA _50_. The well depth *u*_*R*_ for the binding of antigen receptors R and antigens A in the effective double-well adhesion potential is proportional to the antigen concentration A (see eq. (9)). The full data points are determined from eq. (13), the open data points from the approximative eq. (14). The full and dashed lines are interpolation of these data points. As a function of *l*_*C*_ *l*_*R*_, the activation threshold value *u*^50^ is symmetric around the minimum at Δ*l* = |*l*_*C*_ − *l*_*R*_| = 0 and increases up to Δ*l* = Δ*l*^*^ ≃ 8.4 nm at which the antigen-receptor and adhesion-receptor complexes start to segregate. For |Δ*l*| > Δ*l*^*^, activation requires the value [RA]_50_ = *u*_*c*_ = 0.15*k*_*B*_*T a*^2^, which is independent of Δ*l* and of [RA]_50_, because antigen-receptor complexes can only form in segregated domains in which the concentration RA is equal to the concentration [CL] in the domains of the adhesion-receptor complexes CL, as illustrated in (d). (b) For Δ*l* = 0, the adhesion-receptor complexes stabilize adhesion domains with a local membrane separation that is ideal for the formation of antigen-receptor complexes. (c) For 0 < Δ*l* < Δ*l*^*^, the antigen-receptor complexes can form next to adhesion-receptor complexes, but are strained because of the length mismatch.

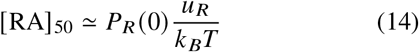

in which the effect of *u*_*R*_ on *P*_*R*_ is neglected. The binding probability *P*_*R*_ 0 is determined from the probability distribution *P*(*l*) for *u*_*R*_ = 0, i.e. from the blue distribution *P*(*l*) in Fig. 2(d), which is independent of Δ*l* = |*l*_*C*_ − *l*_*R*_|. As a function of *l*_*C*_ − *l*_*R*_, the antigen threshold 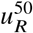 for activation is symmetric around the minimum located at Δ*l* = 0, at which the antigen-receptor and adhesion-receptor complexes have the same length, as depicted in Fig. 3(b). For small to moderate length mismatch Δ*l*, the activation threshold 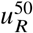 increases with Δ*l* because of the decrease of *P*_*R*_ with Δ*l* (see Fig. 2(b)), which needs to be counterbalanced by larger values of *u*_*R*_ to reach [RA]_50_ in eq. (13). For length mismatches Δ*l* larger than the critical value Δ*l*^*^, the antigen-receptor and adhesion-receptor complexes segregate as depicted in 3(d). This segregation implies that antigen-receptor complexes can only form in domains with a local concentration RA that is equal to the concentration CL in the adhesion-receptor domains (27), which requires 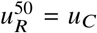, irrespective of Δ*l*. The activation threshold 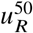 therefore does not further increase for Δ*l* > Δ*l*^*^ and is independent of RA _50_ as long as the complex concentrations RA = CL in the co-existing domains are larger than RA _50_. For small to moderate length mismatches Δ*l* below the critical value Δ*l*^*^ for segregation, in contrast, the antigen-receptor complexes still can form next to adhesion-receptor complexes, but are strained by the length mismatch as depicted in Fig. 3(c), which leads to a decrease of their binding free energy and affinity that needs to compensated by larger antigen concentrations A, i.e. by larger values of *u*_*R*_ in our effective model.

### Modeling of experimental data for the cooperativity of antigen receptors and CD2 adhesion receptors in CAR-T cell activation

To determine how CD2 elongation affects the antigen sensitivity of the TCR and the CARs of Fig. 1, Burton et al. (15) generated Jurkat T cells that express one of these antigen receptors and either wildtype CD2 or an elongated CD2 variant. The CD2 adhesion receptors were elongated by inserting segments of the extracellular (EC) domain of the glycoprotein mucin CD43, which adopts an extended conformation that lacks tertiary structure due to heavy glycosylation. Electron microscopy images of mucin CD43 indicate that each amino acid contributes on average 0.2 nm to the EC domain length (38). A short flexible linker between CD2 and CD43 with amino acid sequence GSSSS was estimated to contribute additional 0.85 nm in length based on a worm-like-chain model and a persistence length of 0.38 nm (15). Therefore, producing a synthetic elongated CD2 molecule with, for example, 40 amino acids from CD43, can be estimated to extend the EC domain length 14.0 nm (39) of the wildtype CD2-CD58 adhesion-receptor complex by 8.85 nm. In the experiments of Burton et al. (15), U87 cells were used as antigen-presenting-cells pulsed with different concentrations of antigenic peptides before being co-cultured with Jurkat T cells expressing each antigen receptor and the different CD2 elongations. The EC50 was determined as the concentration of peptide that lead to half-maximal activation of T cells.

The data points in Fig. 4 represent mean EC50 values of 3 to 6 independent experiments, and the error bars the standard deviation of these independent measurements. The EC50 values of the TCR and STAR are minimal for the length 14 nm of the wildtype CD2 complex and increase with elongation of the CD2 complex up to 19 nm (see Fig. 4(a)). For the CD2 complex lengths of 23 nm and larger, the EC50 values are about two orders of magnitude larger and rather independent of the CD2 length. The EC50 dependence on CD2 complex length thus resembles the functional dependence of the right arm in Fig. 3(a). The EC50 values of the CARs with CD8a, CD28, or IgG1 hinge, in contrast, are minimal for elongated CD2 complexes.

**Figure 4:**
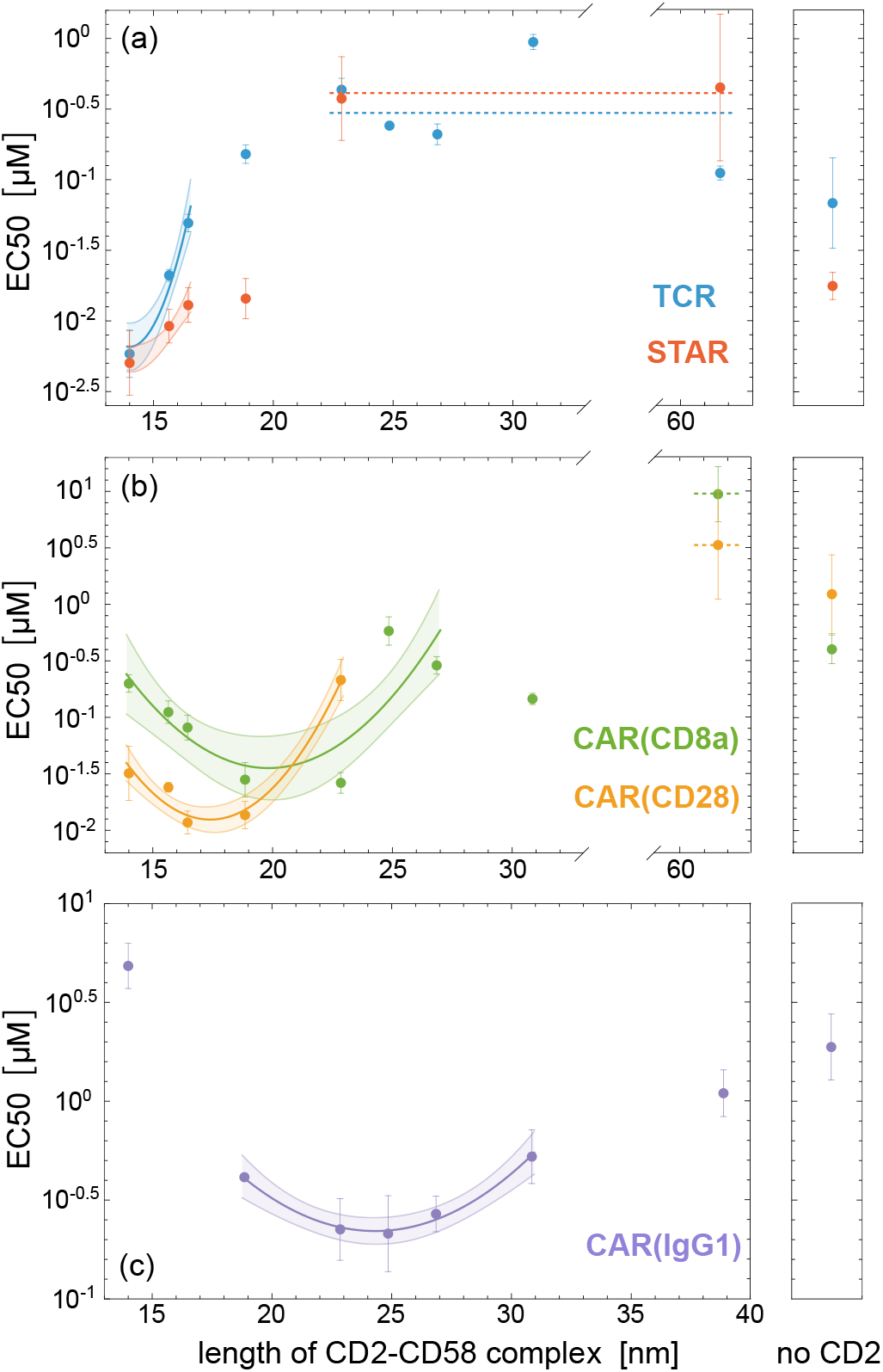
Concentration EC50 of soluble antigenic peptides for half-maximal activation of Jurkat T cells that express the indicated antigen receptor (see Fig. 1) and wildtype or elongated CD2 versus estimated length of the CD2-CD58 complexes. The data points with errors represent the mean and standard deviation of EC50 values obtained from 3 to 6 independent experiments (15). EC50 values of Jurkat T cells that only express the antigen receptor, but not CD2, are shown for comparison. The lines and shaded areas represent fits of eq. (6) with 66% confidence bands to the data points with small to moderate length deviation of the CD2 complex length from the optimal length at which EC50 is minimal. The values for the antigen-receptor complex length *l*_*R*_ and the effective standard deviation ξ_ef_ of function EC50 (*l*_*C*_) obtained from the fits are shown in Table 1.

In our modeling of the experimental data, we fit eq. (6) to the data points for small to moderate length difference from the optimal CD2 complex length at which the antigen concentration EC50 for half-maximal activation is minimal. Eq. (6) is based on the assumption that the distribution *P*(*l*) of the local membrane separation is dominated by the CD2-CD58 complexes for small to moderate length difference *l*_*C*_ − *l*_*R*_ between antigen-receptor and adhesion-receptor complexes. This assumption corresponds to the approximation of eq. (14) in our coarse-grained simulation model, which leads to the open data points in Fig. 3(a). Two central fit parameters in our modelling are the length *l*_*R*_ of the antigen-receptor complex and the effective standard deviation 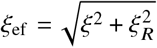, which determine the center and width of the function EC50(*l*_*C*_) (see eq. (6)). The parameter ξ_ef_ depends on the standard deviation ξ of the distribution *P*(*l*) of the local membrane separation and on the standard deviation ξ_*R*_ of the separation-dependent binding constant *k*(*l*) of the antigen-receptor complexes (see eqs. (2) and (3)). The obtained values of the fit parameters are shown in Table 1. The length *l*_*R*_ of the TCR and STAR receptor complex, which cannot be fitted to the data in 4(a), is assumed to be *l*_*R*_ = 14 nm and, thus, equal to the estimated length of the wildtype CD2-CD58 complex. The lengths of the CAR complexes obtained from our fits agree with fit results of Burton et al. (15) within error bounds.

**Table 1:**
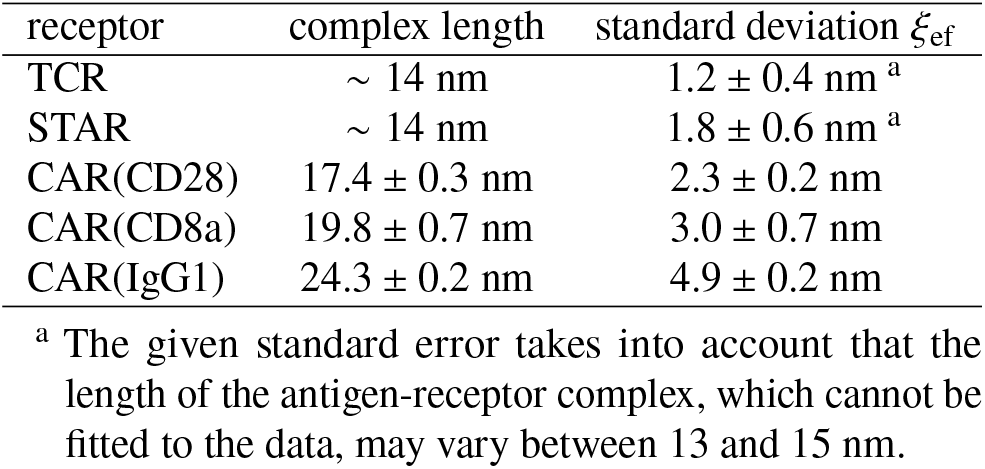
Length *l*_*R*_ of antigen-receptor complexes and effective standard deviation ξ_ef_ of the half-maximally activating antigen concentration EC50 as a function of the adhesion-receptor complex length *l*_*C*_ obtained from the fits in Fig. 4 with eq. (6). The indicated errors are standard errors.

| receptor | complex length | standard deviation $\xi_{\text{ef}}$ |
| --- | --- | --- |
| TCR | $\sim 14$ nm | $1.2 \pm 0.4$ nm<sup>a</sup> |
| STAR | $\sim 14$ nm | $1.8 \pm 0.6$ nm<sup>a</sup> |
| CAR(CD28) | $17.4 \pm 0.3$ nm | $2.3 \pm 0.2$ nm |
| CAR(CD8a) | $19.8 \pm 0.7$ nm | $3.0 \pm 0.7$ nm |
| CAR(IgG1) | $24.3 \pm 0.2$ nm | $4.9 \pm 0.2$ nm |

<sup>a</sup> The given standard error takes into account that the length of the antigen-receptor complex, which cannot be fitted to the data, may vary between 13 and 15 nm.

Burton et al. also determined EC50 values of Jurkat T cells that express the antigen receptor, but no CD2 adhesion receptors. For all antigen receptors, the EC50 values of these CD2-free Jurkat T cells are clearly larger the EC50 values of Jurkat T cells that contain CD2 adhesion receptors of optimal length but also clearly smaller than the EC50 values of Jurkat T cells with a large length mismatch between antigen-receptor and adhesion-receptor complexes (see Fig. 4).

## DISCUSSION

### Cooperativity of antigen-receptor and adhesion-receptor complexes at small to moderate length mismatch

To understand our fit results for the effective standard deviation ξ_ef_ of the functions EC50(*l*_*C*_) in Fig. 4, we first turn to the relative roughness ξ of the opposing membranes. For small to moderate length mismatch Δ*l* = |*l*_*C*_ − *l*_*R*_| of antigen-receptor and adhesion-receptor complexes, the relative membrane roughness ξ is determined by the adhesion-receptor complexes in our model, because the concentration of these complexes then greatly exceeds the concentration of antigen-receptor complexes required for activation. Statistical-physical theory and coarse-grained simulations indicate the relation (17)

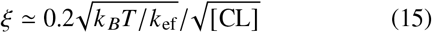

between ξ and the complex concentration [CL]. The numerical factor 0.2 in this relation depends rather weakly on the length and flexibility of the complexes (17) and, thus, should be rather independent of CD2 elongation. The increase of the fit parameter 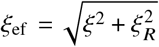 with the length of the antigen-receptor complexes in Table 1 therefore likely result from an increase of the standard deviation ξ_*R*_ of the separation-dependent binding constant *k l* of the antigen-receptor complexes (see eq. (3)). Because ξ is smaller than ξ_ef_ by definition, the relative membrane roughness can be estimated to be about 1 nm to be consistent also with the fit values of ξ_ef_ for TCR and STAR. According to eq. 15, the adhesion-receptor complex concentration [CL] then needs to be larger than 1000 µm^2^ in close-contact zones stabilized by the complexes, because the effective membrane rigidity *κ*_ef_ = *κ*_1_*κ*_2_/(*κ*_1_ + *κ*_2_) varies between 10 and 20 *k*_*B*_*T* for typical cell membrane rigidities *κ*_1_ and *κ*_2_ between 20 and 40 *k*_*B*_*T* (37). For Jurkat T cells adhering to supported bilayers that contain the ligand CD58 of CD2, fluorescence measurements indicate average CD2-CD58 complex concentrations up to 760/µm^2^ in the cell adhesion zones (40). In the cell-cell adhesion experiments of Burton et al. (15), locally larger concentrations of CD2-CD58 complexes exceeding 1000/µm^2^ may result from the observed segregation of large glycocalyx components or from cytoskeletal forces on close contact zones that lead to a compression of the zones.

For the TCR, we obtain ξ_*R*_ ≃ 0.7 nm for the value ξ_ef_ ≃ 1.2 nm and ξ_*R*_ ≃ 1.25 nm for the upper standard-error bound ξ_ef_ ≃ 1.6 nm in Table 1 with the estimated relative membrane roughness ξ ≃ 1 nm. The standard deviation ξ_*R*_ of the separation-dependent binding constant *k l* of the TCR-pMHC complex is likely dominated by the tilting of the EC domain relative to the membranes. For the unbound membrane-anchored TCR with associated CD3 domains, atomistic simulations indicate a tilt angle of about 33^°^ for the TCR EC domain with a standard deviation of about 9^°^ (41). For the TCR-pMHC complex, tilt angle variations with a standard deviation of 9^°^ would lead to ξ_*R*_ ≃ 1.5 nm. However, the membrane anchoring of the pMHC ligand not included in the simulations may reduce the tilt angle variations and lead to values in agreement with the estimates above. For the STAR with a structure similar to the TCR (see Fig. 1), the ξ_ef_ value obtained from our fits is somewhat larger but agrees within standard errors with the value for the TCR. In general, a somewhat larger ξ_*R*_ value of the STAR may result from an effectively more flexible connection of the antigen-binding scFv domain, compared to the TCR. With the estimated relative membrane roughness ξ 1 ≃ nm, we obtain ξ_*R*_ = 1.5 ± 0.7 nm for the STAR.

For the CARs, we obtain ξ_*R*_ = 2.1 ± 0.2 nm for CAR(CD28), ξ_*R*_ = 2.8 ± 0.7 nm for CAR(CD8a), and ξ_*R*_ = 4.8 ± 0.2 nm for CAR(IgG1) from the fit values of ξ_ef_ in Table 1with the estimated relative membrane roughness ξ ≃ 1 nm. The ξ_*R*_ values of the CARs thus increase with the length of the CAR complexes, which indicates an increase in the flexibility of the complexes. For CAR(IgG1), however, the maximal and minimal values of EC50 in Fig. 4(c) differ by only about 1 order of magnitude. The simulation results in Fig. 3(a) indicate that the modeling assumption of a distribution *P l* of the local membrane separation *l* that is dominated by the adhesion-receptor complexes then no longer holds and leads to overestimates of ξ_ef_ in fits based on this assumption. The open data points in Fig. 3(a) obtained with this modeling assumption overestimate 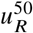 compared to the numerically exact full data points for the exemplary green curves with a difference of about 1 order of magnitude between maximal and minimal 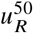 values. The 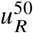 values in Fig. 3(a) are proportional to the antigen concentration and, thus, correspond to the EC50 values of the experiments.

### Segregation of antigen-receptor and adhesion-receptor complexes at large length mismatch

For sufficiently large length mismatch Δ*l*, the antigen-receptor and adhesion-receptor complexes segregate in our coarse-grained model. This segregation implies that the antigen-receptor complexes can only form in domains with a local concentration [RA] that is equal to the concentration [CL] in the domains of the adhesion-receptor complexes (27). In our model, the antigen activation threshold does not further increase for Δ*l* values beyond the critical value Δ*l*^*^ for segregation, because the complex concentrations [RA] = [CL] in the co-existing domains are larger than the concentration [RA]_50_ for half-maximal activation (see Fig. 3(a)). In line with these modelling results, the EC50 values of the TCR and STAR at estimated CD2-CD58 complex lengths of 23 nm and larger tend to be constant (see dashed lines in Fig. 4(a)). For the TCR, the experimentally determined EC50 value for the CD-CD58 complex length of 19 nm is already close to the values for longer CD2-CD58 complexes. Based on a comparison with the expected functional shapes in Fig. 3 obtained in our coarse-grained model, we estimate Δ*l*^*^ ≃ 5 nm for the TCR and Δ*l*^*^ ≃ 7 to 9 nm for the STAR as critical length mismatch for segregation from the CD2-CD58 complexes in the experiments. Our modelling of the experimental data in Fig. 4 thus allows to estimate critical length mismatches for the segregation of TCR and STAR complexes from CD2 complexes at the complex concentrations of the experiments, which we estimate to be of the order of 1000/µm^2^ (see previous section). The obtained critical length mismatches below 10 nm are currently difficult to measure more directly from differences in the membrane separation of segregated domains accessible in high-resolution microscopy experiments of T cells adhering to supported membranes (42). In experiments with synthetic binding proteins anchored to reconstituted lipid vesicles, exclusion of non-binding proteins from vesicle-vesicle adhesion zones has been observed for length differences of about 5 nm between the EC domains of the binding-protein complexes and the non-binding proteins at complex concentrations of several 1000/µm^2^ (43).

The segregation of receptor complexes of different length results from an interplay between the entropy of mixing and the curvature-mediated repulsion of the complexes. This curvature-mediated repulsion increases both with the length mismatch and the concentration of the complexes, because the complexes tend to be closer to each other at higher concentrations in the mixed state, which implies larger membrane curvatures between complexes of different length and, thus, larger membrane bending energies. The general statistical-physical relations summarized in Appendix A allow the prediction of how the onset of segregation depends on the length mismatch Δ*l*, on the concentrations of the complexes, and on the effective bending rigidity *κ*_ef_ of the apposing cell membranes (21, 26, 44). The numerical prefactors in these relations have been determined from simulations with the same coarse-grained model as in the simulations presented here. In this coarse-grained model, an antigen receptor binds to an apposing antigen ligand if the local membrane separation *l* deviates from the complex length *l*_*R*_ by not more than 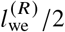, where 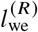 is the width of the binding well. This square-well interaction implies the the separation-dependent binding constant *k* (*l*) is a box function with the value *k* = *k*_*R*_ for 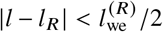 and the value 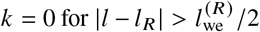. To relate this box function to the Gaussian function *k* (*l*) of Eq. (3) with maximal value *k*_max_, which is used in our modeling of the experimental data in Fig. 4, we note that the box function with the same maximal value *k*_*R*_ = *k*_max_ as the Gaussian function has about the same area for 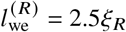. According to this area comparison, the well width 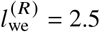 nm of our simulations, for example, corresponds to ξ_*R*_ ≃ 1 nm.

In our coarse-grained model, segregation of the complexes occurs if the width 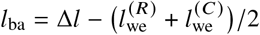 of the barrier between the two binding wells for the antigen-receptor and adhesion-receptor complexes exceeds a threshold value (see Appendix A). This barrier width depends both on the length mismatch Δ*l* and on the binding well widths 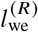 and 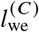 of the antigen-receptor and adhesion-receptor complexes. The standard deviations ξ_*R*_ = 0.7±0.5 nm and ξ_*R*_ = 1.5±0.7 nm for the separation-dependent binding constant *k*(*l*) of the TCR and STAR estimated in the Results section lead to the estimates 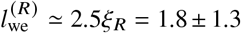 nm for the TCR and 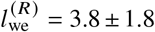 for the STAR. For the exemplary value *l* ^(*C*)^ ≃ 2.5 nm and the estimated critical length mismatches Δ*l*^*^ ≃ 5 nm for the TCR and Δ*l*^*^ ≃ 7 to 9 nm for the STAR (see above), we obtain the estimated critical barrier widths 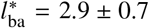 nm for the TCR and 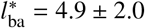 nm for the STAR, which agree within the estimation errors. The larger value of ξ_*R*_ estimated for the STAR thus helps to understand the larger critical value Δ*l*^*^ of the length mismatch, compared to the TCR. With these estimates for 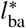 and 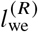, we obtain estimated complex concentrations of the order of 1000 µm^2^ in the segregated domains from eq. (21) for effective rigidity values *κ*_ef_ between 10 and 20 *k*_*B*_*T*, which agree with the complex concentrations estimated based on eq. (15) above. For CAR(CD28) and CAR(CD8a), the critical length mismatch Δ*l*^*^ for segregation can be estimated to be larger than 10 nm based on the fits in Fig. 4(b), in line with the larger estimated values of ξ_*R*_ for these CARs, compared to the TCR and STAR. In addition, the longer CD2 complexes optimal for CAR activation are likely more flexible than the wildtype CD2 complexes, which leads to larger values of 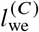.

## CONCLUSION

In this article, we have presented a statistical-physical theory for the membrane-mediated cooperativity of antigen receptors and adhesion receptors in CAR-T cell activation. Our theory quantifies how the effective two-dimensional binding constant *K* of antigen-receptor complexes depends on small to moderate length mismatches to adhesion-receptor complexes and on the thermal membrane roughness ξ (see eq. (4)), which in turn depends on the concentration of the abundant adhesion-receptor complexes that hold the membranes together against thermal membrane shape fluctuations. A central molecular length scale in this theory is the standard deviation ξ_*R*_ of the membrane-separation-dependent binding constant *k l* of the antigen-receptor complexes (see eq. (1)), which reflects the conformational and tilting flexibility of the complexes. Our fits of experimental data for the length-mismatch-dependent antigen sensitivity of CAR-T cells allow to estimate a thermal membrane roughness ξ of about 1 nm in close-contact adhesion zones and complex concentrations of the order of 1000/µm^2^ of CD2-CD58 adhesion-receptor complexes in these zones. According to our fits, ξ_*R*_ varies from about 1 nm for TCR and STAR to values between 2 and 3 nm for the longer CAR(CD8a) and CAR(IgG1).

In addition, we have performed coarse-grained simulations to corroborate our statistical-physical model and to explore the antigen sensitivity in the full range of length mismatches between antigen-receptor and adhesion-receptor complexes. In our coarse-grained simulation model, we have chosen an adhesion-receptor complex concentration of about 800 µm^2^ and a value ξ_*R*_ of about 1 nm, which is within the parameter ranges obtained from the fits of our statistical-physical model to experimental data. Beyond the critical length mismatch of about 8.4 nm for the chosen parameter values, the antigen-receptor and adhesion-receptor complexes segregate, and the antigen concentration required for activation in the simulation model becomes independent of the length mismatch (see Fig. 3), consistent with experimental observations (see Fig. 4(a)).

## APPENDIX A: STATISTICAL PHYSICS OF LENGTH-DEPENDENT SEGREGATION DURING MEMBRANE ADHESION

Integrating out the degrees of freedom of the antigen receptors, adhesion receptors, and their ligands in our coarse-grained model of cell adhesion leads to an effective model of two homogeneous membranes interacting via double-well potential with well depths *u*_*R*_ and *u*_*c*_ that depend on the concentrations and binding constants of the receptors and ligands. The free energy of membrane segments bound in the antigen-receptor well with width 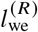 scales as 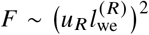, and the free energy of membrane segments bound in the adhesion-receptor well with width 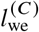 scales as 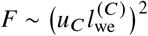 for small values of *u*_*R*_ and *u*_*C*_ (21). These two wells are separated by a barrier of width

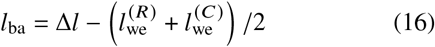

where Δ*l* = *l*_*C*_ − *l*_*R*_ is the length mismatch of the antigen-receptor and adhesion-receptor complexes. Statistical-physical theory indicates that the two membranes exhibit lateral phase separation if the free-energy height of the barrier exceeds a critical value that is inversely proportional to 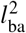 (45).

According to this theory, lateral phase separation into domains of antigen-receptor complexes and domains of adhesion-receptor complexes occurs for

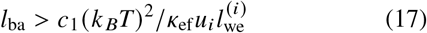

with *i* = R and C and a numerical prefactor *c*_1_ = 0.225 ± 0.02 that has been obtained by fitting MC results for the critical point of phase separation determined with the cumulant method of Fig. 5 at various well widths and barrier widths of the double-well potential (21). For the well width 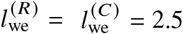 nm, effective bending rigidity *κ*_ef_ = 10 *k*_*B*_*T*, and potential depths *u*_*C*_ = *u*_*R*_ = 0.15 *k*_*B*_*T*/*a*^2^ with *a* = 10 nm, eq. (17) predicts the critical length mismatch Δ*l*^*^ = 8.5 ± 0.5 nm, which agrees with the value Δ*l*^*^ = 8.4 ± 0.1 nm obtained from the MC data in Fig. 5 within the statistical errors.

**Figure 5:**
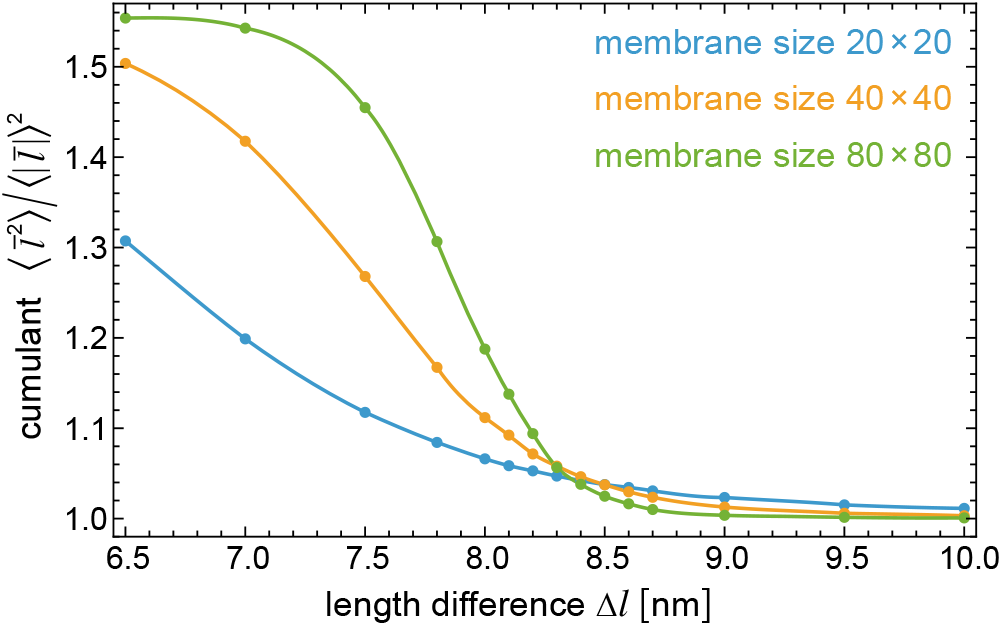
MC results for the cumulant 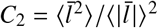 of the spatially averaged membrane separation 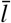 as a function of the length mismatch Δ*l* of the antigen-receptor and adhesion-receptor complexes for membranes with different sizes of 20 × 20, 40 × 40, and 80 × 80 patches and periodic boundaries. In these simulations, the membranes interact via an effective double-well potential *V*_ef_ with equal well depths *u*_*R*_ = *u*_*C*_ = 0.15*k*_*B*_*T a*^2^ where *a* = 10*nm* is the side length of the patches, well width *l*_we_ = 2.5 nm, and a distance of Δ*l* between the centers of the wells (see Fig. (2(b)). The data points represent MD data with statistical errors that are smaller than the point sizes, and the lines are interpolations.

To obtain statistical-physical statements on how the critical length mismatch Δ*l*^*^ depends on the concentrations [RA] and [CO] of the antigen-receptor and adhesion-receptor complexes, we now aim to replace the effective potential depths *u*_*R*_ and *u*_*C*_ in eq. (17) by the complex concentrations. The complex concentrations [RA] = *P*_*R*_*u*_*R*_/*k*_*B*_*T* and [CO] = *P*_*C*_*u*_*C*_/*k*_*B*_*T* depend on the probabilities *P*_*R*_ and *P*_*C*_ that the membranes are bound in the antigen-receptor and CD2 adhesion-receptor well, respectively, which in turn depend on well depths *u*_*R*_ and *u*_*C*_ (see Methods). For membrane domains bound in one of the wells, statistical-physical theory and MC data lead to (26)

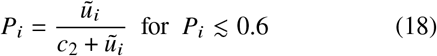

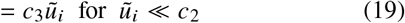

with = *R* or *C*, numerical factors *c*_2_= 0.071 ± 0.002 and *c*_3_ = 13 ± 1, and rescaled potential depths 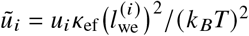. According to eqs. (17) and (19), lateral segregation of the complexes occurs for (36)

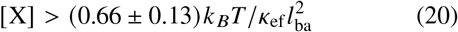

with [X] = [CO] = [RA] . Eq. (19), however, does not hold for the rescaled potential depth *ũ*_*C*_ = 0.09375 of the coarse-grained model considered here, or in other words, for complex concentrations of the order of 1000 µm^2^ estimated based on the experimental data of Fig. 4 (see Discussion). From eqs. (17) and (18), we obtain the more general condition for lateral segregation with 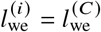 or 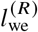 depending on the complex, and numerical factors *c*_1_ and *c*_2_ defined above.

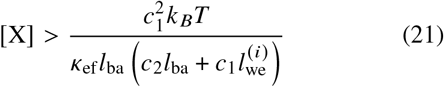

## APPENDIX B: FINITE-SIZE METHOD FOR DETERMINING THE CRITICAL POINT OF LENGTH-BASED SEGREGATION WITH MC SIMULATIONS

At the critical point for the length-based segregation of receptor-ligand complexes, the correlation length of the membrane shape fluctuations becomes infinite. Following standard finite-size simulation methods for determining critical points (46), we determine the cumulant 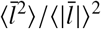 of the spatially averaged membrane separation 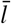 with MC simulations for membranes of different sizes. Here, ⟨…⟩ denotes temporal averages over the MC simulations. In our simulations, the membranes interact via an effective double-well potential with equal well depths of 0.15*k*_*B*_*T*/*a*^2^ where *a* = 10 nm is the side length of the patches, well width *l*_we_ = 2.5 nm, and distance Δ*l* between the centers of the wells. The cumulant decreases from the value π/2 ≃ 1.57 for small Δ*l*, at which the distribution *P*(*l*) of the local membrane separation is bimodal (see e.g. the purple distribution *P l* in Fig. 2(d)) to the value 1 for large Δ*l*, with a functional form that depends on the membrane size (46). At large Δ*l*, the distribution *P*(*l*) obtained from our simulations is unimodal and centered in only one of the wells. At the critical length mismatch Δ*l*^*^ for segregation of the complexes, the value of the cumulant *C*_2_ becomes independent of the membrane size because of the infinite correlation length of membrane shape fluctuations at this critical point (21, 46). From the intersection points of the interpolation lines in Fig. 5 at Δ*l* = 8.49 nm for the sizes 20 × 20 and 40 × 40, Δ*l* = 8.36 nm for the sizes 20 × 20 and 80 × 80, and Δ*l* = 8.29 nm for the sizes 40 × 40 and 80 × 80, we obtain Δ*l*^*^ = 8.4 ± 0.1 nm. The MC data have been determined in simulations with a length of up to 10^10^ MC steps per patch. In a single MC step of our simulations, the local separation *l*_*i*_ of two apposing membrane patches *i* is attempted to be shifted to a new value 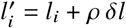 where *ρ* is a random number between -1 and 1 and *δl* is the step width. The MC step is accepted or rejected based on the standard Metropolis criterion, and the step width *δl* is adjusted during the simulation such that the average acceptance ratio of MC steps is 0.4 (44).

## AUTHOR CONTRIBUTIONS

All authors designed the research. T.R.W. carried out all simulations and calculations. T.R.W. and O.D. wrote the article. All authors edited the article.

## ACKNOWLEDGMENTS

T.R.W. is deeply grateful to Wolfgang Helfrich for setting him on a scientific path as a diploma student, and thanks the Max Planck Society for generous financial support.

## REFERENCES

1. Rossjohn, J., S. Gras, J. J. Miles, S. J. Turner, D. I. Godfrey, and J. McCluskey, 2015. T cell antigen receptor recognition of antigen-presenting molecules. Annu Rev Immunol 33:169–200.

2. Labanieh, L., and C. L. Mackall, 2023. CAR immune cells: design principles, resistance and the next generation. Nature 2023 614:7949 614:635–648.

3. Minguet, S., M. V. Maus, and W. W. Schamel, 2025. From TCR fundamental research to innovative chimeric antigen receptor design. Nat. Rev. Immunol. 25:212–224.

4. Siller-Farfan, J. A., and O. Dushek, 2018. Molecular mechanisms of T cell sensitivity to antigen. Immunol. Rev. 285:194–205.

5. Irvine, D. J., M. A. Purbhoo, M. Krogsgaard, and M. M. Davis, 2002. Direct observation of ligand recognition by T cells. Nature 419:845–849.

6. Huang, J., M. Brameshuber, X. Zeng, J. Xie, Q.-j. Li, Y.-h. Chien, S. Valitutti, and M. M. Davis, 2013. A Single Peptide-Major Histocompatibility Complex Ligand Triggers Digital Cytokine Secretion in CD4(T Cells. Immunity 39:846–857.

7. Harris, D. T., M. V. Hager, S. N. Smith, Q. Cai, J. D. Stone, P. Kruger, M. Lever, O. Dushek, T. M. Schmitt, P. D. Greenberg, and D. M. Kranz, 2018. Comparison of T cell activities mediated by human TCRs and CARs that use the same recognition comains. J Immunol 200:1088– 1100.

8. Gudipati, V., J. Rydzek, I. Doel-Perez, V. D. R. Gonçalves, L. Scharf, S. Königsberger, E. Lobner, R. Kunert, H. Einsele, H. Stockinger, M. Hudecek, and J. B. Huppa, 2020. Inefficient CAR-proximal signaling blunts antigen sensitivity. Nat. Immunol. 21:848–856.

9. Salter, A. I., A. Rajan, J. J. Kennedy, R. G. Ivey, S. A. Shelby, I. Leung, M. L. Templeton, V. Muhunthan, V. Voillet, D. Sommermeyer, J. R. Whiteaker, R. Gottardo, S. L. Veatch, A. G. Paulovich, and S. R. Riddell, 2021. Comparative analysis of TCR and CAR signaling informs CAR designs with superior antigen sensitivity and in vivo function. Sci. Signal. 14:eabe2606.

10. Burton, J., J.A. Siller-Farfán, J. Pettmann, B. Salzer, M. Kutuzov, P. A. van der Merwe, and O. Dushek, 2023. Inefficient exploitation of accessory receptors reduces the sensitivity of chimeric antigen receptors. Proc. Natl. Acad. Sci. USA 120:e2216352120.

11. Patel, A., V. Andre, S. B. Eguiguren, M. I. Barton, J. Burton, E. M. Denham, J. Pettmann, A. M. Mørch, M. A. Kutuzov, J.A. Siller-Farfán, M. L. Dustin, P. A. van der Merwe, and O. Dushek, 2024. Using CombiCells, a platform for titration and combinatorial display of cell surface ligands, to study T-cell antigen sensitivity modulation by accessory receptors. EMBO J. 43:132–150.

12. Majzner, R. G., and C. L. Mackall, 2018. Tumor antigen escape from CAR T-cell therapy. Cancer Discovery 8:1219–26.

13. Davis, S. J., S. Ikemizu, M. K. Wild, and P. a. van der Merwe, 1998. CD2 and the nature of protein interactions mediating cell-cell recognition. Immunological reviews 163:217–36.

14. Sušac, L., M. T. Vuong, C. Thomas, S. von Bülow, C. O’Brien-Ball, A. M. Santos, R. A. Fernandes, G. Hummer, R. Tampé, and S. J. Davis, 2022. Structure of a fully assembled tumor-specific T cell receptor ligated by pMHC. Cell 185:3201–3213.

15. Burton, J., J.A. Siller-Farfán, V. Andre, E. Jenkins, M. I. Barton, S. Bustamante Eguiguren, J. Cabezas Caballero, S. J. Davis, T. R. Weikl, P. A. van der Merwe, and O. Dushek, 2025. Optimising CAR-T cell sensitivity by engineering matched extracellular sizes between CAR/antigen and CD2/CD58 adhesion complexes. bioRxiv https://www.biorxiv.org/content/early/2025/01/06/2025.01.06.631424.

16. Pettmann, J., A. Huhn, E. A. Shah, M. A. Kutuzov, D. B. Wilson, M. L. Dustin, S. J. Davis, P. A. van der Merwe, and O. Dushek, 2021. The discriminatory power of the t cell receptor. eLife 10:1–42.

17. Xu, G.-K., J. Hu, R. Lipowsky, and T. R. Weikl, 2015. Binding constants of membrane-anchored receptors and ligands: A general theory corroborated Monte Carlo simulations. J. Chem. Phys. 143:243136.

18. Hu, J., G.-K. Xu, R. Lipowsky, and T. R. Weikl, 2015. Binding kinetics of membrane-anchored receptors and ligands: Molecular dynamics simulations and theory. J. Chem. Phys 143:243137.

19. Steinkühler, J., B. Rozycki, C. Alvey, R. Lipowsky, T. R. Weikl, R. Dimova, and D. E. Discher, 2019. Membrane fluctuations and acidosis regulate cooperative binding of ‘marker of self’ protein CD47 with the macrophage checkpoint receptor SIRP??. J. Cell. Sci. 132:jcs216770.

20. Weikl, T. R., and R. Lipowsky, 2001. Adhesion-induced phase behavior of multicomponent membranes. Phys. Rev. E. 64:011903.

21. Asfaw, M., B. Rozycki, R. Lipowsky, and T. R. Weikl, 2006. Membrane adhesion via competing receptor/ligand bonds. Europhys. Lett. 76:703–709.

22. Lipowsky, R., 1996. Adhesion of Membranes via Anchored Stickers. Phys. Rev. Lett. 77:1652–1655.

23. Weikl, T. R., and R. Lipowsky, 2004. Pattern formation during T-cell adhesion. Biophys. J. 87:3665–3678.

24. Tsourkas, P. K., N. Baumgarth, S. I. Simon, and S. Raychaudhuri, 2007. Mechanisms of B-cell synapse formation predicted by Monte Carlo simulation. Biophys. J. 92:4196–4208.

25. Reister-Gottfried, E., K. Sengupta, B. Lorz, E. Sackmann, U. Seifert, and A. S. Smith, 2008. Dynamics of specific vesicle-substrate adhesion: From local events to global dynamics. Phys. Rev. Lett. 101:208103.

26. Krobath, H., B. Rozycki, R. Lipowsky, and T. R. Weikl, 2009. Binding cooperativity of membrane adhesion receptors. Soft Matter 5:3354–3361.

27. Różycki, B., R. Lipowsky, and T. R. Weikl, 2010. Segregation of receptor–ligand complexes in cell adhesion zones: phase diagrams and the role of thermal membrane roughness. New J. Phys. 12:095003.

28. Weil, N., and O. Farago, 2010. Entropy-driven aggregation of adhesion sites of supported membranes. Eur. Phys. J. E 33:81–87.

29. Krobath, H., B. Rozycki, R. Lipowsky, and T. R. Weikl, 2011. Line tension and stability of domains in celladhesion zones mediated by long and short receptorligand complexes. PLoS One 6:e23284.

30. Bihr, T., U. Seifert, and A.-S. Smith, 2012. Nucleation of ligand-receptor domains in membrane adhesion. Phys. Rev. Lett. 109:258101.

31. Li, L., J. Hu, X. Shi, Y. Shao, and F. Song, 2017. Lipid rafts enhance the binding constant of membrane-anchored receptors and ligands. Soft Matter 13:4294–4304.

32. Knezevic, M., H. Jiang, and S. Wang, 2018. Active Tuning of Synaptic Patterns Enhances Immune Discrimination. Phys. Rev. Lett. 121:238101.

33. Li, L., J. Hu, L. Li, and F. Song, 2019. Binding constant of membrane-anchored receptors and ligands that induce membrane curvatures. Soft Matter 15:3507–3514.

34. Helfrich, W., 1973. Elastic properties of lipid bilayers: Theory and possible experiments. Z. Naturforsch. C 28:693–703.

35. Lipowsky, R., 1988. Lines of renormalization group fixed points for fluid and crystalline membranes. Europhys. Lett. 7:255–261.

36. Weikl, T. R., 2018. Membrane-mediated cooperativity of proteins. Annu. Rev. Phys. Chem. 69:521–539.

37. Steinkühler, J., E. Sezgin, I. Urbancic, C. Eggeling, and R. Dimova, 2019. Mechanical properties of plasma membrane vesicles correlate with lipid order, viscosity and cell density. Commun. Biol. 2:337.

38. Cyster, J. G., D. M. Shotton, and A. F. Williams, 1991. The dimensions of the T lymphocyte glycoprotein leukosialin and identification of linear protein epitopes that can be modified by glycosylation. EMBO J. 10:893–902.

39. Wang, J. H., A. Smolyar, K. Tan, J. H. Liu, M. Kim, Z. Y. Sun, G. Wagner, and E. L. Reinherz, 1999. Structure of a heterophilic adhesion complex between the human CD2 and CD58 (LFA-3) counterreceptors. Cell 97:791–803.

40. Zhu, D.-M., M. L. Dustin, C. W. Cairo, and D. E. Golan, 2007. Analysis of two-dimensional dissociation constant of laterally mobile cell adhesion molecules. Biophys. J. 92:1022–1034.

41. Pandey, P. R., B. Różycki, R. Lipowsky, and T. R. Weikl, 2021. Structural variability and concerted motions of the T cell receptor – CD3 complex. eLife 10:e67195.

42. Velas, L., M. Brameshuber, J. B. Huppa, E. Kurz, M. L. Dustin, P. Zelger, A. Jesacher, and G. J. Schütz, 2021. Three-Dimensional single molecule localization microscopy reveals the topography of the immunological synapse at isotropic precision below 15 nm. Nano Lett2. 21:9247–9255.

43. Schmid, E. M., M. H. Bakalar, K. Choudhuri, J. Weichsel, H. S. Ann, P. L. Geissler, M. L. Dustin, and D. A. Fletcher, 2016. Size-dependent protein segregation at membrane interfaces. Nat. Phys. 12:704.

44. Weikl, T. R., and R. Lipowsky, 2006. Membrane adhesion and domain formation. In Advances in Planar Lipid Bilayers and Liposomes. A. Leitmannova Liu, editor. Academic Press.

45. Lipowsky, R., 1994. Discontinuous unbinding transitions of flexible membranes. J. Phys. II 4:1755–1762.

46. Ammann, A., and R. Lipowsky, 1996. Discontinuous phase transitions of membranes: a Monte Carlo study. J3. Phys. II 6:255–270.

